# Segmentation-less, automated vascular vectorization robustly extracts neurovascular network statistics from in vivo two-photon images

**DOI:** 10.1101/2020.06.15.151076

**Authors:** Samuel A. Mihelic, William A. Sikora, Ahmed M. Hassan, Michael R. Williamson, Theresa A. Jones, Andrew K. Dunn

## Abstract

Recent advances in two-photon microscopy (2PM) have allowed large scale imaging and analysis of blood vessel networks in living mice. However, extracting a network graph and vector representations for vessels remain bottlenecks in many applications. Vascular vectorization is algorithmically difficult because blood vessels have many shapes and sizes, the samples are often unevenly illuminated, and large image volumes are required to achieve good statistical power. State-of-the-art, three-dimensional, vascular vectorization approaches often require a segmented (binary) image, relying on manual or supervised-machine annotation. Therefore, voxel-by-voxel image segmentation is biased by the human annotator or trainer. Furthermore, segmented images oftentimes require remedial morphological filtering before skeletonization or vectorization. To address these limitations, we present a vectorization method to extract vascular objects directly from unsegmented images without the need for machine learning or training. The Segmentation-Less, Automated, Vascular Vectorization (SLAVV) source code in MATLAB is openly available on GitHub. This novel method uses simple models of vascular anatomy, efficient linear filtering, and low-complexity vector extraction algorithms to remove the image segmentation requirement, replacing it with manual or automated vector classification. SLAVV is demonstrated on three in vivo 2PM image volumes of microvascular networks (capillaries, arterioles and venules) in the mouse cortex. Vectorization performance is proven robust to the choice of plasma- or endothelial-labeled contrast, and processing costs are shown to scale with input image volume. Fully-automated SLAVV performance is evaluated on simulated 2PM images of varying quality all based on the large (1.4×0.9×0.6 mm^3^ and 1.6×10^8^ voxel) input image. Vascular statistics of interest (e.g. volume fraction, surface area density) calculated from automatically vectorized images show greater robustness to image quality than those calculated from intensity-thresholded images.

**Author summary:** Samuel Mihelic is a PhD candidate in the Biomedical Engineering Department at the University of Texas at Austin. He graduated from Oregon State University (Chemical Engineering BS, Mathematics BS). He hosts the GitHub repository for the code used in this article: https://github.com/UTFOIL/Vectorization-Public. His research interests are in-vivo neural microvascular image analysis, anatomy, and plasticity.

William Sikora graduated with a BS in Computational Biomedical Engineering from The University of Texas at Austin in May 2020. He is working with Dr. Yuan Yang and the Laureate Institute for Brain Research as a PhD student of Biomedical Engineering at the University of Oklahoma in Tulsa, researching the highly non-linear world of neural coupling and its link to common neurological pathologies such as stroke.

Ahmed Hassan is a graduate of the University of California, Los Angeles and the University of Texas at Austin with a Bachelor's degree in Microbiology, Immunology, and Molecular Genetics and an MSE/PhD in Biomedical Engineering. His graduate research was concentrated in imaging and instrumentation, and his interests include developing optical and laser systems for neuroimaging, image processing and reconstruction, and advanced image analysis.

Michael Williamson earned a BSc (Honours) in Neuroscience in 2016 from the University of Alberta, where he trained with Dr. Fred Colbourne. He is currently a doctoral student at the University of Texas at Austin working in the labs of Drs. Theresa Jones and Michael Drew.

Theresa Jones is a Professor in the Department of Psychology and Neuroscience at The University of Texas at Austin. Her laboratory studies plasticity of neural structure and synaptic connectivity following brain damage and injury.

Andrew K. Dunn is the Donald J. Douglass Centennial Professor of Engineering in the Department of Biomedical Engineering at The University of Texas at Austin and the Director of the Center for Emerging Imaging Technologies. His research focuses on the development of innovative optical imaging techniques for studying the brain.

## Introduction

The neurovascular system provides oxygen and nutrients in response to local metabolic demands through the process of neurovascular coupling [1]. This process is dysregulated in pathological conditions such as hypertension and Alzheimer’s disease [2, 3]. Individual capillary tracking over multiple imaging sessions would provide a useful experimental tool for measuring neurovascular plasticity in preclinical disease models. Such experiments could screen new stroke therapeutics and provide much needed relief to clinical studies [4–7]. However these measurements remain intractable for large image volumes, due to difficulties in vectorization.

A vectorized network consists of a graph that summarizes the connectivity and a collection of simple objects that represent individual vessel segments. High-fidelity vectorization is difficult to attain in general, but greatly facilitates and simplifies the vascular network for statistical analysis [8] or blood flow simulation [9]. The state of the art approach to vascular vectorization is skeletonization, which requires a nearly-perfect image segmentation of the vascular network. From a computer vision perspective, the segmentation of blood vessels from in vivo optical microscopy images presents major challenges:

- Blood vessels have many sizes and bifurcations causing a variety of shapes.
- Objects are unevenly illuminated (especially larger vessels), and image quality decreases with depth.
- Large, information-rich images are required to achieve both statistical power and spatial resolution

It is this image segmentation requirement, which is difficult to meet in general, that motivates us to revisit the vascular vectorization workflow.

For the purposes of quantifying vascular anatomy, skeletonization techniques are used to vectorize and extract the centerlines of blood vessels [10–13]. Skeletonization techniques perform iterative morphological filtering on binarized images, and therefore require segmented images generated from vessel enhancement filtering, thresholding, or manual or machine-learned annotation. Manual, voxel-by-voxel, image segmentation ensures a high-quality binary input image to skeletonize, but is a tedious and often heuristic task. Researchers demonstrated individual capillary tracking in living mouse cortex over several weeks, but estimates of vascular plasticity lacked statistical significance due to the small, manually segmented, sample size [14]. Alternatively, convolutional neural networks allow computers to learn this manual task from example [15, 16]. However, deep learners that are trained by humans in voxel-by-voxel classification have human biases and are not intrinsically robust to input image properties such as resolution and noise level.

Many filters using local curvature information show improved robustness to image quality [17–19]. These filters require eigenvalue decomposition of the Hessian at many voxels, and are thus computationally expensive. They rely on local shape information, are difficult to extend to multiscale, and show attenuated response at vessel bifurcations. These deficiencies were addressed by [20] using a unitless ratio of eigenvalues. [21] used a Hessian-based filter to segment images using an exploratory vectorization algorithm that automatically traces vessels and terminates each trace when the filtered image drops below a predefined threshold. However, the termination criteria was arbitrary and did not produce a connected network.

Manual or machine-learning approaches to voxel-by-voxel, image segmentation [22] do not guarantee any topological structure without remedial morphological filtering. Furthermore, traditional performance metrics of image segmentation (e.g. Dice coefficient) do not measure topological or connectivity accuracy. As an alternative, we propose extracting vascular vectors directly from 2PM images, thereby enforcing fundamental shape and connectivity constraints. To demonstrate this workflow, we present the Segmentation-Less, Automated, Vascular Vectorization (SLAVV) method outlined in Figure 1.

**Fig. 1.**
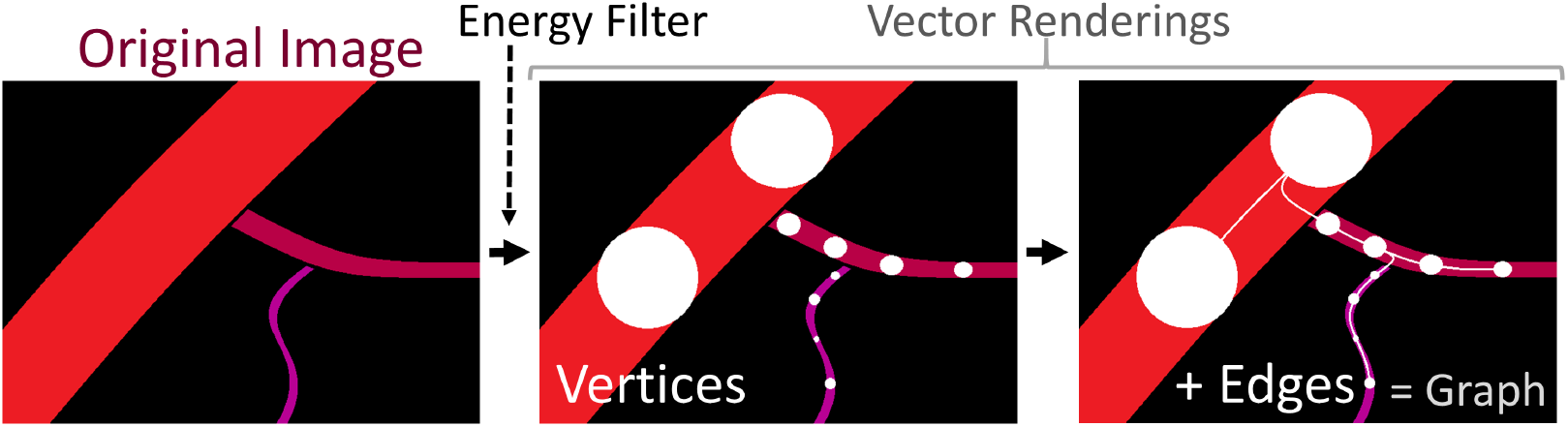
Overview of the SLAVV approach. The purpose of SLAVV is to vectorize vascular objects from raw three dimensional images. The first step of the method is to linearly filter the input image to form “energy” and “size” images, which enhance vessel centerlines and estimate vessel sizes, respectively. Next, vertices along the blood vessels are extracted as local minima of the 3D energy image. Vertices are then connected by edges, which follow minimal energy trajectories. Finally, a graph theoretic representation of the vascular network is generated from the vertices and edges.

The advantage of the direct vectorization approach is that there is no requirement to segment, interpolate, or otherwise preprocess the input image. The image processing required to vectorize is simplified: Vessel segments and bifurcations are both enhanced by a single blob detector. The extracted vectors have realistic shape and connectivity constraints, so there is no need for nonlinear morphological filtering. Additionally, there is no need to create training sets or train the software, because the method does not rely on machine learning. However, the extracted vectors are probabilistic and need to be classified in some way. A graphical user interface is used to curate the extracted vectors from in vivo, plasma- and endothelial-labeled 2PM images to create ground-truth vectorizations. Fully-automated vectorization performance is then evaluated on realistic, simulated 2PM images of varying quality. Performance is evaluated as the percent error in several vascular statistics of interest: volume fraction, surface area density, length density, and bifurcation density.

## Materials and methods

### 2PM Imaging

Two-photon fluorescence microscopy (2PM), in vivo, three-dimensional images of murine microvasculature were acquired from two mice at two resolutions with two different sources of contrast. Table 1 summarizes the images input to the SLAVV software for demonstration.

**Table 1.**
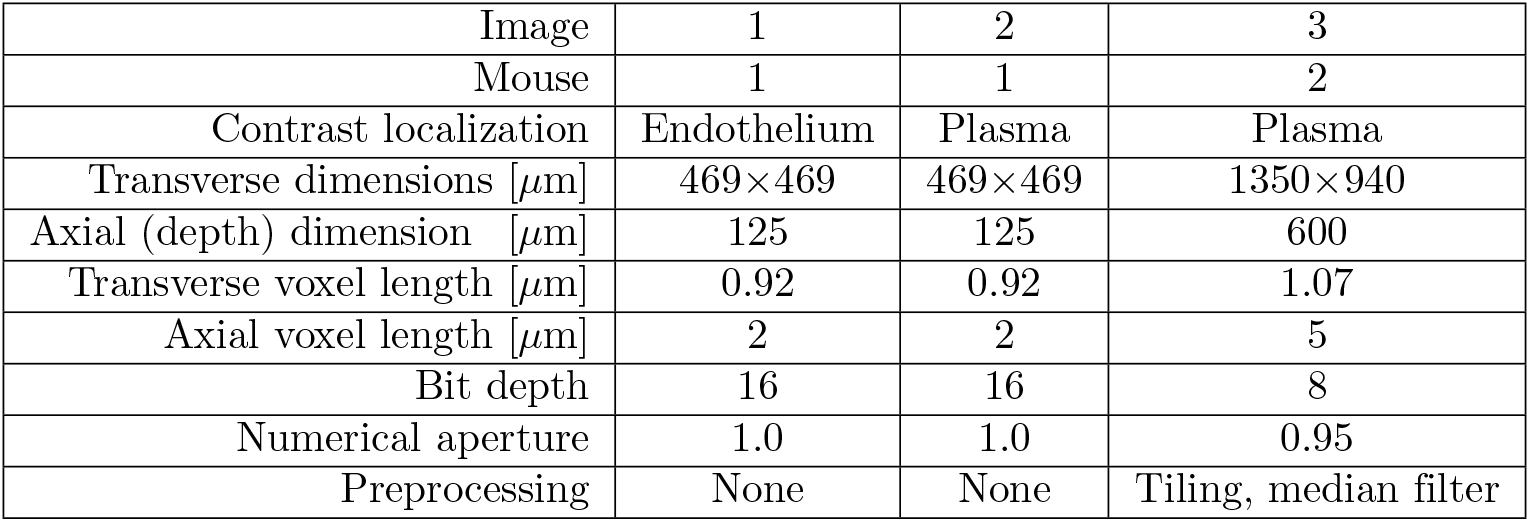
Descriptions of 2PM input images used in SLAVV demonstration. Note that the endothelium (vessel wall) was labeled in Image 1.

| Image | 1 | 2 | 3 |
| --- | --- | --- | --- |
| Mouse | 1 | 1 | 2 |
| Contrast localization | Endothelium | Plasma | Plasma |
| Transverse dimensions [$\mu\text{m}$] | 469×469 | 469×469 | 1350×940 |
| Axial (depth) dimension [$\mu\text{m}$] | 125 | 125 | 600 |
| Transverse voxel length [$\mu\text{m}$] | 0.92 | 0.92 | 1.07 |
| Axial voxel length [$\mu\text{m}$] | 2 | 2 | 5 |
| Bit depth | 16 | 16 | 8 |
| Numerical aperture | 1.0 | 1.0 | 0.95 |
| Preprocessing | None | None | Tiling, median filter |

### Dual channel plasma- and endothelial-labeled

Mouse 1 (Young adult Tie2-GFP, FVB background, JAX stock no. 003658) was implanted with a 4mm cranial window over sensorimotor cortex [23]. For imaging, the mouse was anesthetized with 1-2 % isoflurane in oxygen and head fixed. A retro-orbital injection of 0.05mL of 1 % (w/v) 70 kDa Texas Red-conjugated dextran was given to label the plasma. GFP fluorescence was localized to the endothelium. Images 1 and 2 were acquired with a Prairie Ultima two-photon microscope with a Ti:Sapphire laser (Mai Tai, Spectra-Physics) tuned to 870 nm and a 20× 1.0 NA water immersion objective (Olympus) using Prairie View software.

### Large volume tiling

Mouse 2 was injected intravenously with Texas Red and imaged through a cranial window using a two-photon microscope [24]. The laser (1050 nm, 80 MHz repetition rate, 100 fs pulse duration [25]) was scanned using galvo-galvo mirrors over 550 *μ*m × 550 *μ*m (512 × 512 pixels) and axially using a motorized stage over 600 *μ*m depth (120 slices). The images were output as 16 bit signed integers (3×10^7^ voxels). To achieve the tiling in Image 3, a 2×3 grid of plasma-labeled images with 100 *μ*m overlaps were median-filtered (3×3×3 voxel kernel) and then tiled with translational registration via cross-correlation [26] in Fiji [27].

### Automated vessel vectorization

Novel Segmentation-Less Automated Vascular Vectorization (SLAVV) software is developed in MATLAB to extract vector sets representing vascular networks from raw gray-scale images without the need for pre-processing or specialized hardware. The SLAVV method has four major steps:

1. Energy: multi-scale linear filtering
2. Vertex extraction
3. Edge extraction
4. Network and strand identification

Vertices and edges resulting from Steps 2 and 3 are probabilistic vectors that require classification (see Vector classification).

### Energy: multi-scale linear filtering

The raw (unprocessed/uninterpolated) input 3D image is matched-filtered for vessels (idealized as spherical objects) within a user-specified size range to yield a multi-scale, 4D image (Figure 2, conceptual). The matched filter is the convolution of a Laplacian of Gaussian (LoG) with standard deviation, *σ*, and an Ideal (spherical and/or annular) kernel of radius *r*. Therefore, the radius, *R*, of the matched vessel is given by *R^2^* = *σ^2^* + *r^2^*.

**Fig. 2.**
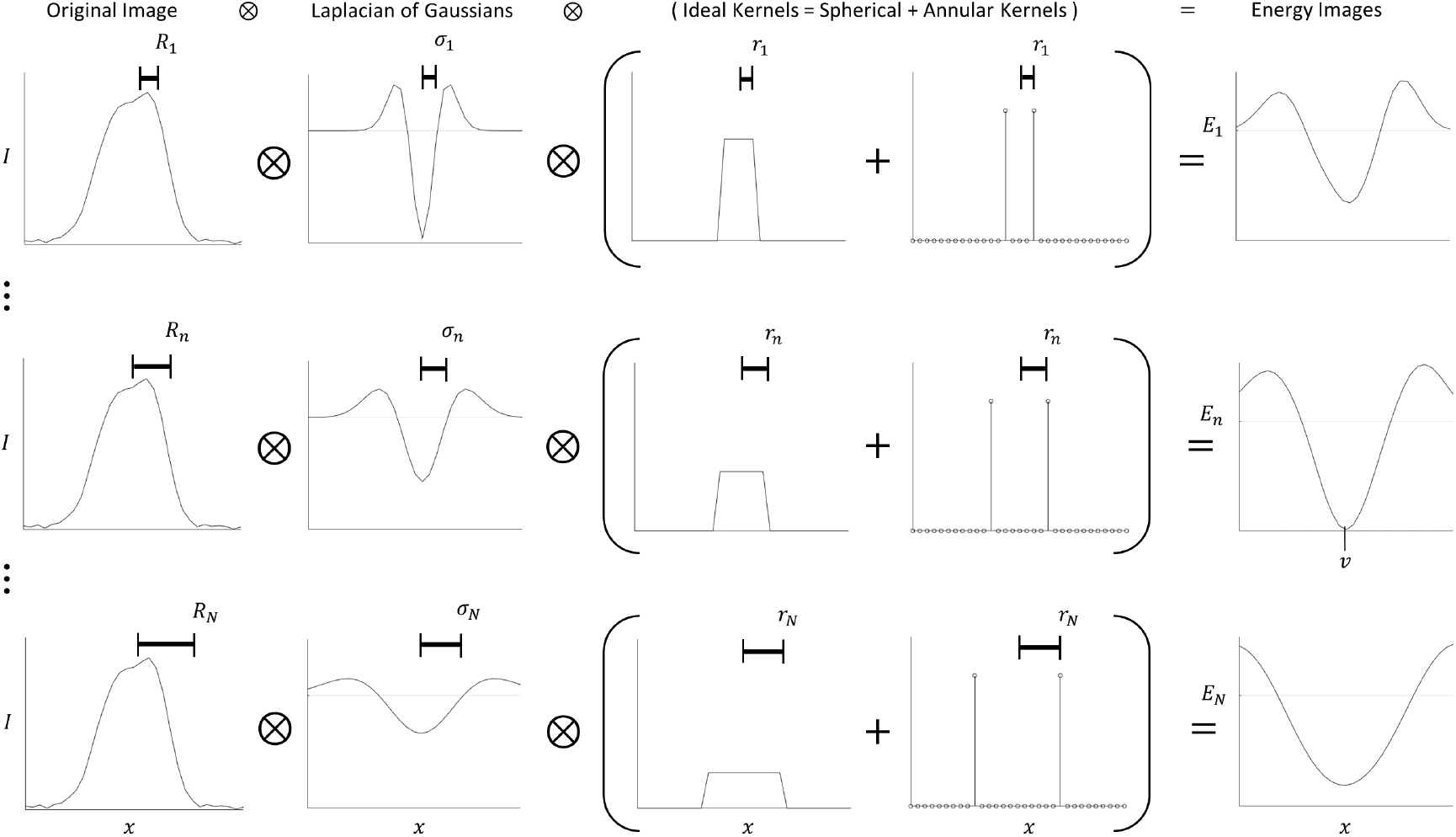
One-dimensional simplification of linear filtering step. To form the energy image *E* at scales 1*…n…N*, the original image *I*(*x*) is convolved with a LoG filter and an Ideal kernel. The Ideal kernel is a linear combination of spherical and annular pulses to match the fluorescent signal shape of vessels. *σ^2^* is the variance of the Gaussian, *r* is the radius of the Ideal kernel, so *R^2^* = *σ^2^* + *r^2^* is the square “radius” of the LoG, matched filter. The resulting multiscale energy image is projected along the scale coordinate to form two 3D images that depict energy and size (not shown here, example in Figure 3). In this example, the kernel weighting factor was chosen so that the sums of all the spherical and annular kernels were all equal, the ratio *r/σ* was chosen to be 1, and a vertex was found at location *v* with radius *R_n_* and energy *E_n_*(*v*).

To detect a variety of fluorescent signal shapes, the Ideal kernel is a combination of spherical and annular pulses. The weighting parameter *f_S_* controls the fraction of spherical versus annular pulse in the matching kernel, thus weighting the responses from signals in the shape of the vessel wall vs lumen. Therefore, *f_S_* = |*K_S_*|/(|*K_S_*| + |*K_A_*|), where |*K_S_*| and |*K_A_*| are the total weights (*L^1^* norms) of the spherical and annular matching kernels.

The Ideal kernel is convolved with a Gaussian to achieve robustness to noise and slight mismatches in size and shape. The processing parameter *f_G_* controls the relative sizes of the Gaussian and Ideal kernels, thereby trading noise robustness for accuracy of size and position. Therefore, *f_G_* = *σ/*(*σ* + *r*), where *σ* is the standard deviation of the Gaussian kernel and *r* is the radius of the Ideal kernel.

The scale space sampling is exponentially distributed with tunable density. The octave is defined as doubling of the vessel volume. For example, Image 1 is sampled across 16 doublings of the matched filter volume at a rate of 6 scales per octave, producing 96 discrete scale samples. To enforce a minimal amount of blurring and ensure good shape agreement between objects in the image and matched filters, a 3D (anisotropic) Gaussian model of the microscope PSF [28] is also convolved with each matched filter. Table 2 shows the processing parameters used to process the featured images.

**Table 2.** Processing parameters used to vectorize the three experimental images and the set of synthetic images derived from Image 3.

| Image | 1 | 2 | 3 | 3-synthetic |
| --- | --- | --- | --- | --- |
| $f_G$ [%] | 50 | 75 | 50 | 60, 80, 100 |
| $f_S$ [%] | 50 | 50 | 85 | 100 |
| Smallest radius [$\mu\text{m}$] | 1.5 | 1.5 | 1.5 | 1 |
| Largest radius [$\mu\text{m}$] | 60 | 60 | 60 | 45 |
| Scales per octave | 6 | 6 | 3 | 3 |

The 4D (multi-scale) energy image is projected across the scale coordinate to yield two 3D images (each with the same dimensions as the input) called the energy and size images. The energy image encodes “energy” which is used to estimate the likelihood of that voxel containing a vessel centerline. The size image encodes the expected radius of a vessel centered at each voxel. Because energy is the Laplacian of a real-valued image, negative values correspond to locally bright regions of the original image. Image voxels with positive energies are therefore ignored based on the assumption that the fluorescent signal is brighter than the background.

To compare Laplacian values across different scales and amounts of blurring, each second derivative is weighted by the variance of the Gaussian filter at that scale in that dimension [29] to compensate for decreased derivative values due to blurring. The projection method from multi-scale, 4D image back to 3D depends on the signal type, because the annular signal is more sensitive to size mismatches and cannot be averaged across many sizes reliably. Thus, for the annular input signal (endothelial label), the estimated scale is simply the scale that yielded the least energy. For the spherical input signal (plasma label), the estimated scale is a weighted average across all scales that yielded a negative energy using energy magnitude as weights. For linear combinations of annular and spherical signals, the weighted average of the two estimates is taken. The 3D energy image is then defined by looking up the 4D energy image at the scale nearest the estimated scale at each voxel.

To improve computational efficiency, several design features are built into the energy filtering step. For larger scale objects, the image is downsampled in each dimension before filtering to have a resolution no greater than 10 voxels per object radius, and then upsampled afterwards with linear interpolation. The image volume is processed in overlapping chunks to respect memory constraints and allow for parallelization. The overlapping length is expanded for larger scales to eliminate edge effects within the image volume. To minimize computational complexity, all blurring, matched filtering, and derivative approximations are calculated in the Fourier domain with a single, combined filter for each scale sampled from analytical Fourier representations.

### Vertex extraction

The purpose of the vertex extraction step is to identify centerline points along the vascular network to serve as seed points for the edge extraction step. The inputs are the energy and size images, and the output is a set of vertices: non-overlapping spheres that should be concentric with vessel centerlines, share the same radius as the vessel at that central location, and densely mark the vasculature with at least one vertex for each strand in the final output network.

This algorithm is based on the keypoint extraction outlined in [29]: Vertex center points are located by searching the energy image for local minima in all three-dimensions. Additionally, the sizes of the vertices are assigned by referencing the size image at their center points. Vertex volumes are then painted onto a blank canvas from lowest to highest energy to obtain the most probable non-overlapping subset (Algorithm 1, 1-dimensional example).

#### Algorithm 1

Extracting vertices from the energy and radius images.

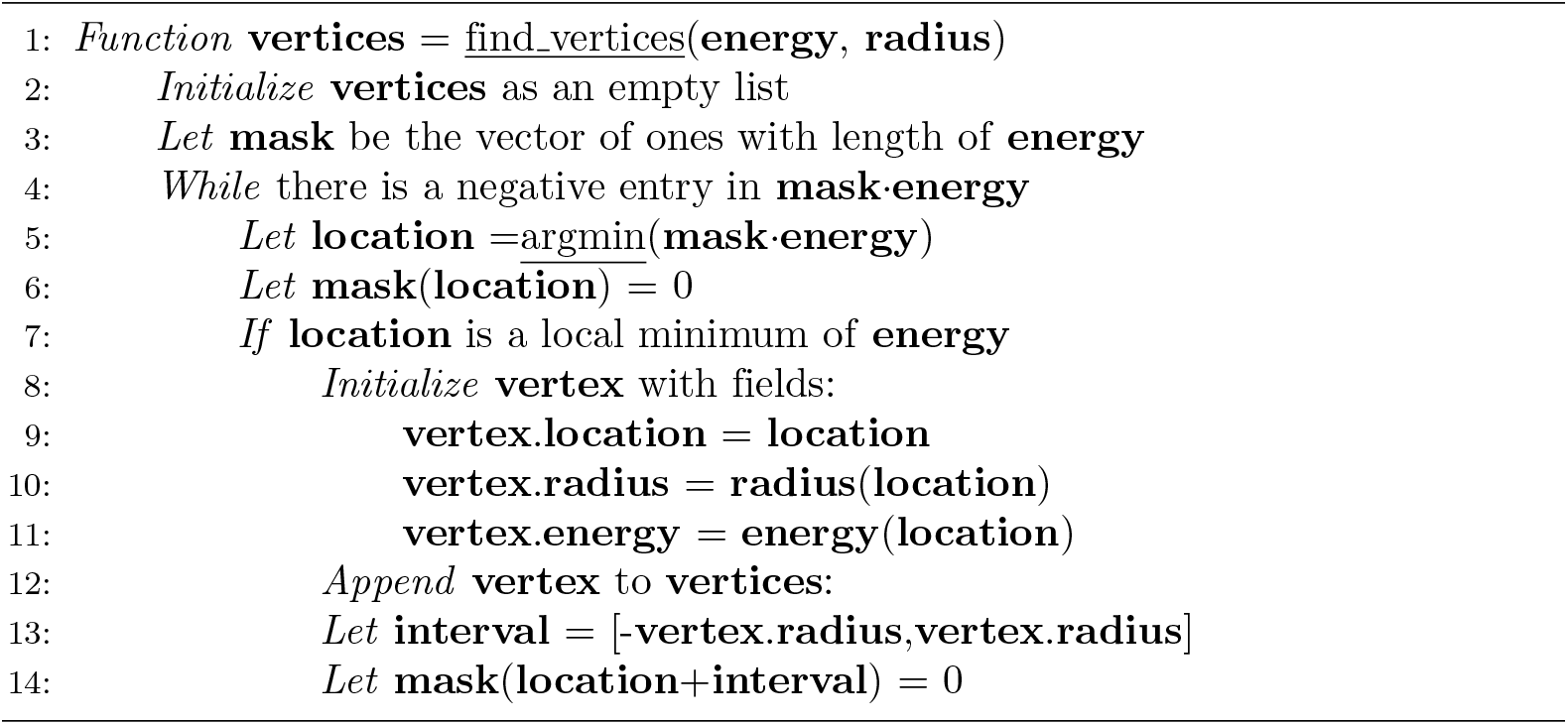

### Edge extraction

The purpose of the edge extraction step is to trace the vessel segments between pairs of vertices with a simple automated algorithm. The inputs are the set of vertices and the energy and size images. The output is a set of edges: vessel traces connecting vertex pairs, summarized by ordered lists of centerpoint 3-space coordinates and associated radii. Edges are ordered from most to least likely by the maximum energies along their traces. For edges with the same energy, the shorter one is considered more likely. This decision selects for the child edge whenever a child and parent have the same maximum energy.

Edges are traced by deterministically exploring the space around each vertex by repeatedly visiting the lowest energy neighbor, in a watershed manner, until a terminal vertex is found (Algorithm 2). When a terminal vertex is explored, the newfound edge is traced back to the original vertex following the shortest path with lowest maximum energy. To guarantee that this algorithm uses finite time and memory, maxima were placed on the number of edges that each seed vertex could find (4 edges per vertex) and on the length of trace (20-100 times seed vertex radius).

#### Algorithm 2

Extracting edges associated to a vertex set from an energy image.

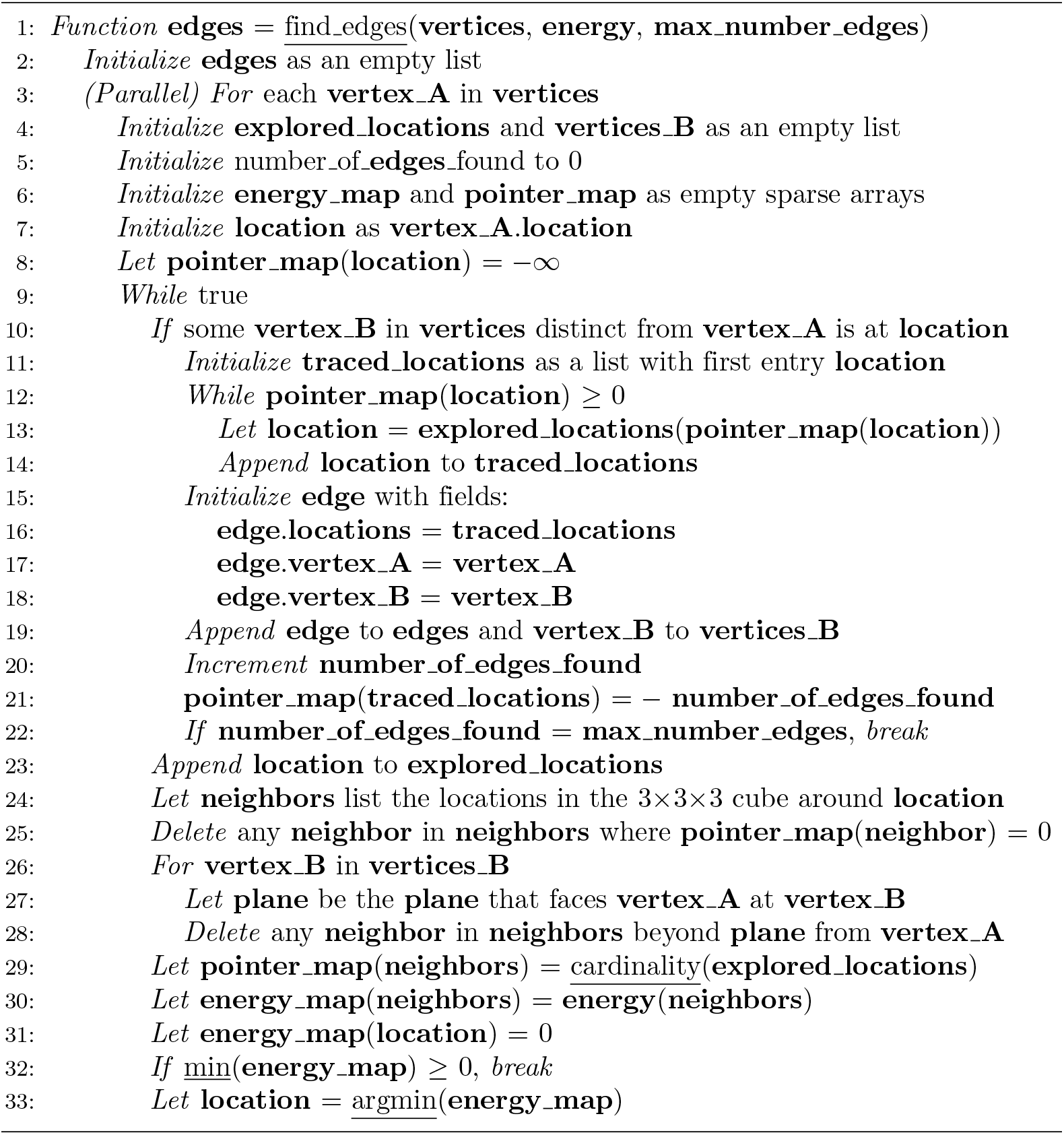

Algorithm 2 extracts edges from each origin vertex in the order of increasing maximum energies. To avoid double-counting stretches of vessel segments, vertices are not allowed to make multiple connections in the same direction. Once a vertex discovers a neighbor vertex by means of an edge, it is not allowed to search farther in the direction of that neighbor.

Algorithm 2 extracts non-overlapping centerline traces for the edges from each origin vertex. When two edge traces with the same origin vertex would share a common segment of centerline voxels, the algorithm only counts the redundant trace in the first-extracted, lower-energy (more-probable) edge. That edge is called the parent, and the higher-energy one the child edge. To promote vertex placement near vessel bifurcations, vertices are added (after Algorithm 2) wherever a child edge meets its parent.

In cases where two vertices discover each other, only the most probable edge that connects them is kept. Small cycles of three fully-connected vertices are eliminated using graph theoretic methods. Two vertices are adjacent by a small cycle if they can be mutually reached using one and two edges. The connected components of the adjacency matrix by these small cycles is calculated, and the least probable edge is removed from each component. The adjacency matrix is then recalculated and the removal repeated until there are no adjacencies by small cycles.

### Network and strand identification

The purpose of the network and strand identification step is to organize the vectors to facilitate statistical calculations and visualization. The inputs are the sets of vertex and edge objects, and the outputs are sets of strands, bifurcations and endpoints. Bifurcations and endpoints are special vertices: bifurcations connect three or more edges, endpoints connect one. A strand is a set of one or more consecutive edges that connects exactly two special vertices.

Bifurcations are found by calculating the adjacency matrix for the graph of the vertices and edges. Bifurcations are vertices associated with 3 or more edges. These vertices and their associated edges are then removed from the graph, leaving only vertices with 1 or 2 edges. The connected components of this graph identify the majority of the strand objects. The remaining strands and strand fragments are the single edges that connect to bifurcation vertices.

Once all the edges are assigned to strands and a subset of the vertices to bifurcations, a smoothing operation is applied that keeps the special vertices fixed and smooths the positions and sizes along the one-dimensional strands. Gaussian smoothing kernels are defined for discrete locations along each strand with standard deviations equal to their radii. Smoothing kernels are further weighted by the energy value at each strand location. Algorithm 2 guarantees negative energy at every location along an edge, so the weighting is well defined and will favor the more likely, lower-energy locations. The variable smoothing kernel is then applied to each strand location to spatially average along the strand trace to increase the statistical significance of the five local quantities: 3-position, size, and energy.

## Vector classification

Due to the property of the SLAVV method extracting probabilistic vectors directly from grayscale images (without the need for image segmentation), a vector classification step after the edge extraction step is used to provide a deterministic input to the network statistic calculations. Vector classification is performed manually or automatically.

### Interactive curation software

Two built-in, graphical curator interfaces are used to manually classify probabilistic vertices and edges into true and false categories. The human effort spent on each manual curation is described here and automatically recorded by the curation software.

The automatically generated vectors are rendered transparently over the raw image. The user sets the intensity limits of the underlying raw image to accommodate variable brightness and contrast. The user may also view the probabilistic vectors with their brightness dependent on their energies to assess the accuracy of the automated filtering and vector extraction steps. The interface enables navigation within the input image volume to view any rectangular sub-volume as a maximum intensity projection in *z*. Manual edits to vectors can then be made within the sub-volume.

Manual edits are either classifications or additions. Two vector classification tools are used: point-and-click true/false toggling and local energy thresholding (within the navigation volume). Some vertices are added to a volume in a manual way that is assisted by the energy and size images. The user specifies the *x* and *y* location with a point-and-click and the software automatically selects the *z* position that minimizes the energy and the size from the corresponding location in the size image. Additionally, some edges are added by selecting two vertices to connect. The location and size along the added edge are simply linearly interpolated from the two vertices.

### Automated energy thresholding

The vertex and edge objects are both segmented using global thresholds on the energy feature. Receiver operating characteristic (ROC) curves are created by sweeping this energy threshold. The most accurate operating points are chosen with the knowledge of the ground truth image to maximize voxel-by-voxel classification accuracy. For the vertex objects, global thresholding applies a user-defined upper limit on the acceptable energy associated with any vertex. Vertices with energies above the threshold are classified as false and eliminated. Since each vertex serves as a seed point (and possible termination point) in the edge extraction algorithm, vertices are segmented prior to edge extraction to improve performance. For the edge objects, global thresholding applies a user-defined upper limit on the acceptable maximum energy associated with any edge. Edges with maximum energy above the threshold are eliminated.

## Visualizations

### Three-dimensional scalar fields

Outputs of the energy, vertex, edge, and network steps of the SLAVV software (see Automated vessel vectorization) are automatically output as TIFF files of 3D scalar fields at the resolution of the input image. These files are opened in ImageJ [27] for viewing and projection. Vertices and edges are rendered as energy-weighted, centerline- or volume-filled objects. Centerline filling creates single voxel wide, continuous traces. Volume filling places spherical structuring elements of estimated radius, concentric with the centerline traces.

### Projections and perspectives

The strand-resolved and depth-resolved *z*-projections are created in MATLAB by rendering strand centerline voxels with colors based on the unique strand identity or the *z*-component of each centerline location, respectively. Three-dimensional visualization is done in MATLAB with the isosurface and isonormals functions by upsampling the strand sample points, and rendering each as a volume-filled object. The colors are again mapped from the strand unique identifiers.

### Statistical analysis of vectors

Statistical calculations are performed within the SLAVV software to extract features of the network output such as volume, surface area, length, and number of bifurcations. These calculations operate on the network output strands idealized as consecutive circular, right cylinders (Figure 4B). For analyzing distributions (histograms) of vessel quantities, the lateral surface area is used as the weighting quantity, because the lateral surface area is directly proportional to the chemical flux across the vessel wall.

### Simulating 2PM images

Realistic 2PM images of varying contrast and noise are simulated from a ground truth vectorization for the purposes of measuring automated vectorization performance. First the ground truth vectorization is rendered in a binary volume-fill at high resolution, blurred with an overestimated Gaussian model for the PSF [28], and downsampled to the resolution of the original raw image. The intensities are then linearly transformed to have a positive background value *I_B_* and a larger foreground value *I_F_*.

To simulate Poisson distributed noise, a normally distributed random variable is added to each voxel with variance equal to the voxel intensity. The foreground and background values are varied to achieve different contrast and noise levels. Contrast is defined as the difference between the foreground and background intensities, *I_F_* − *I_B_*, and noise as the uncertainty in that difference, 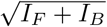. For comparison purposes, simulated image quality is summarized by the contrast to noise ratio.

### Evaluating vectorization performance

The SLAVV method is designed to extract bulk statistics of interest from large volumes of vasculature. Therefore, vectorization performance is evaluated by its accuracy in computing such statistics. The surface area density is a fundamental vascular statistic because it is roughly proportional to the chemical transport per volume of tissue. Other oftentimes reported statistics of interest are the volume fraction, length density, and bifurcation density.

## Results

### Demonstration of SLAVV

To demonstrate the SLAVV approach, Images 1, 2, and 3 were vectorized (see Automated vessel vectorization), manually curated (see Interactive curation software), and visualized (see Visualizations). The runtimes of automated steps are shown in Table 3, and bulk statistics (see Statistical analysis of vectors) are shown in Table 4. Example inputs (plasma- or endothelial-labeled fluorescent images) and intermediates (energy, size, vertex, and edge images) were produced for a small volume of vasculature in Figure 3A. Input images were linearly filtered at many scales (see Energy: multi-scale linear filtering) to produce the energy and size images which estimate the centerline positions and radius of vessels, respectively. Vertices were extracted (see Vertex extraction) as the minima of the energy image, with associated radii looked up from the size image. Edges were traced (see Edge extraction) from one vertex to another along the path of minimum energy, and the size of the vessel at each location was again read from the size image. The connectivity information was then summarized (see Network and strand identification) as a network connecting bifurcations or endpoints with strands. The strand objects (collections of non-bifurcating edges) were each labeled with a unique color in the perspective rendering in Figure 3B. The axial projections in Figure 3C show vector vessel depth or direction information as a volume filling (at one quarter of the estimated vessel radius) overlaying the input image. The runtimes of the automated vectorization steps are shown in Table 3 for each of the three input images. Table 4 shows total network statistics of interest.

**Table 3.** Runtimes of automated processing steps in seconds.

|  | Image 1 | Image 2 | Image 3 |
| --- | --- | --- | --- |
| Energy filter runtime [s] | 986 | 1,037 | 7,853 |
| Vertex extraction runtime [s] | 22 | 35 | 96 |
| Edge extraction runtime [s] | 73 | 204 | 141 |
| Network analysis runtime [s] | 19 | 0.5 | 34 |
| Number of Voxels | $1.67 \times 10^7$ | $1.67 \times 10^7$ | $1.64 \times 10^8$ |

**Table 4.** Bulk network statistics of interest.

|  | Image 1 | Image 2 | Image 3 |
| --- | --- | --- | --- |
| Num. bifurcations | 130 | 194 | 3,383 |
| Length [mm] | 16.2 | 17.0 | 443 |
| Area [mm<sup>2</sup>] | 0.495 | 0.534 | 13.0 |
| Volume [mm<sup>3</sup>] | $2.17 \times 10^{-3}$ | $2.46 \times 10^{-3}$ | $4.40 \times 10^{-2}$ |
| Image Volume [mm<sup>3</sup>] | $2.75 \times 10^{-2}$ | $2.75 \times 10^{-2}$ | 0.761 |

**Fig. 3.**
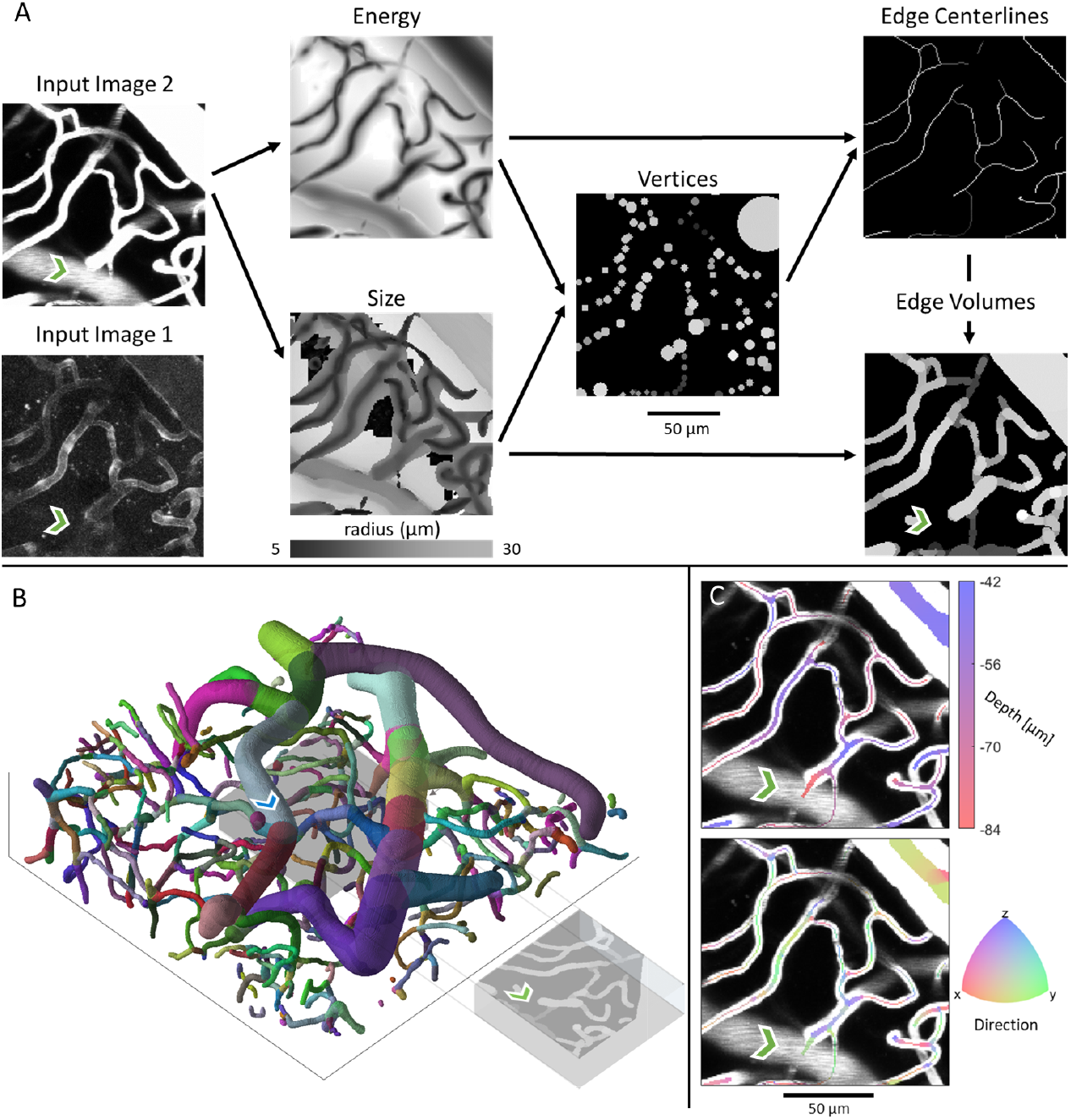
Example projections of original two-photon images, intermediate outputs, and vector renderings of manually assisted SLAVV applied to Image 2. The green chevron is directly underneath a medium sized vessel which bleeds into the original projection volume for the plasma-labeled Image 2 but not the endothelial-labeled Image 1. A. Either Image 1 or 2 could be the input (Image 2 outputs shown). The original image is subject to multiscale, LoG, matched filtering to obtain 3D energy and size images. The energy image is used to estimate vertex centers and the size image to estimate their radii. Vertices are used as genesis and terminus points for energy image exploration in the centerline extraction algorithm. Finally, indexed edge volumes are recalled from the size image to form the volume-filled vector rendering. Gray-scale coloring in the vector renderings corresponds to the energy values and thus vector probabilities. B. 3D visual output of SLAVV. Colors represent strands, which are defined as non-bifurcating vessel segments. Each strand is assigned a random color. The image is 125 *μ*m in *z* and 460 in *x* and *y*. The projection volume used in the other panels is shown as a gray box in the center of the larger volume. The blue chevron marks the vessel that bleeds into Image 2 at the green chevron. C. Depth and direction outputs from SLAVV. Vector volumes are rendered over the original projection at a quarter of their original radius. Direction is calculated by spatial difference quotient with respect to edge trajectory. The centerline for the vessel above the green chevron lies above the projection volume.

**Fig. 4.**
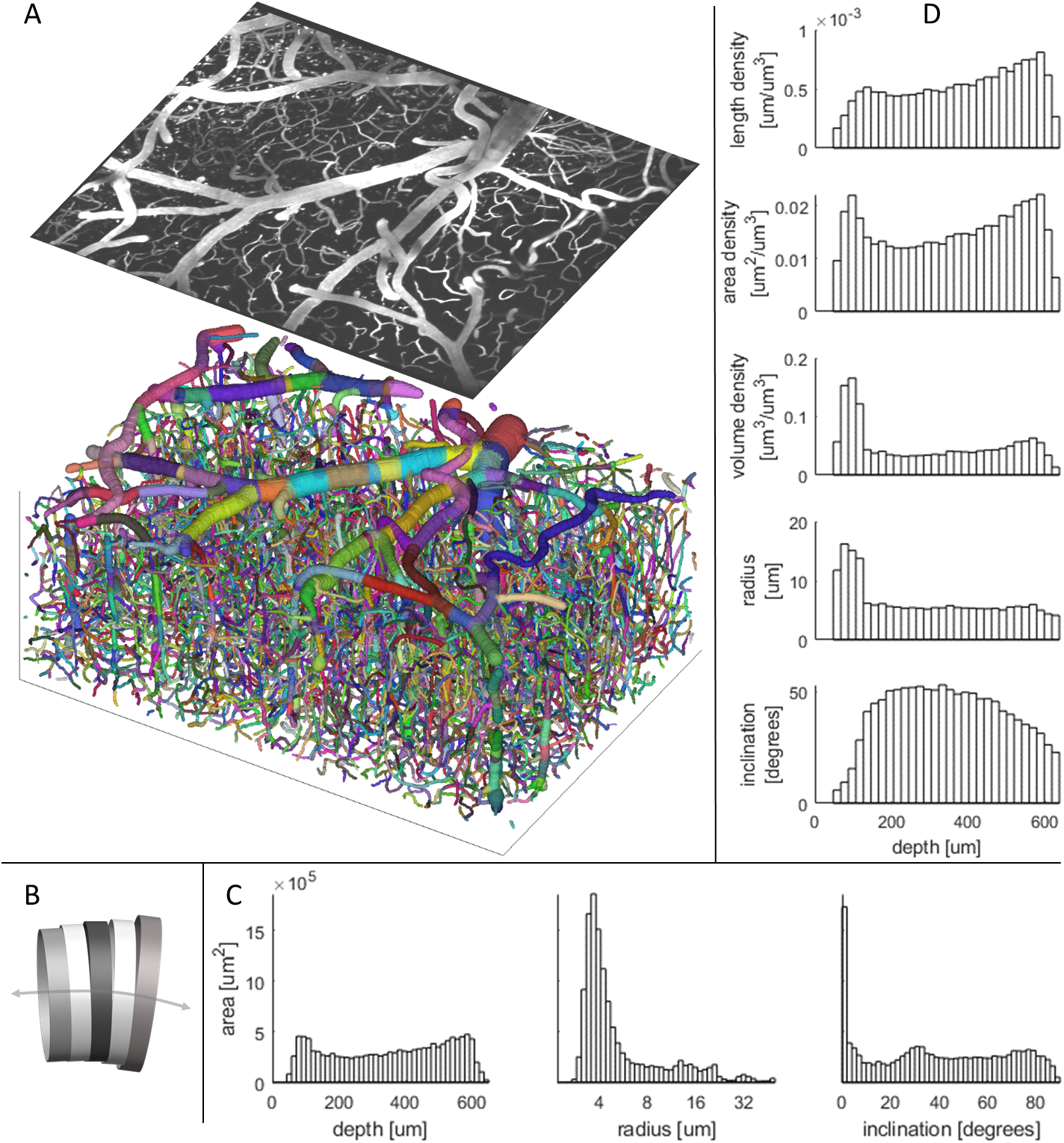
Example statistics of the microvasculature calculated from manually assisted SLAVV applied to Image 3. A. 3D rendering of strand objects similar to Figure 3B. Size of the image is 600 *μ*m in *z*, 1350 in *x*, and 940 *y*. Projection of the first 70 *μ*m of the original image is shown with the same perspective. B. Cartoon depiction of consecutive cylinder representation of a vessel segment used in calculations. C. Histograms of depth, radius, and inclination (angle from the *xy* plane). The large peak in the inclination histogram at horizontal alignment is due to low axial resolution (5 *μ*m). Contributions of cylinders to the bins are weighted by their lateral areas. D. Depth-resolved anatomical statistics output from SLAVV. Cylinders are binned by depth. Their heights, lateral areas, and volumes are summed and divided by the image volume apportioned to each bin to yield length, area, and volume densities of vasculature. Average radius and inclination in each depth bin are lateral area-weighted averages.

### Interactive curation software

For the vectorization of raw input images, an interactive vector curation interface (see Interactive curation software) was used to segment the automatically-generated vertex and edge objects into true and false categories and to patch false negatives. Vertex and edge curation costs for the three curated image volumes are shown in Table 5. Image 3 required the most time because it was the largest volume. Image 1 required more time than Image 2 although it was the same field of view, because the endothelial label was not as strong as the plasma label. The large number of vertex selections for Image 3 were due to the imperfect tiling process (see Large volume tiling) causing image discontinuities away from the image boundaries. Many of these selections were done simultaneously using a click-and-drag box selection. Local thresholds were placed on the energy feature of the vertices of the max energy feature of the edges in order to coarsely classify vectors over large regions of the image at once. The Final Total rows refer to the number of vector objects (vertices or edges) remaining in the final curated datasets.

**Table 5.** Summary of human effort toward manual vector classification on the graphical curator interface in the demonstration of SLAVV. Selections were point-and-click classifications of objects either individually or over a rectangular volume.

|  |  | Image 1 | Image 2 | Image 3 |
| --- | --- | --- | --- | --- |
| Vertices | Duration [min] | 101 | 24 | 206 |
|  | Local Thresholds | 2 | 5 | 110 |
|  | Selections | 4724 | 230 | 19,845 |
|  | Additions | 62 | 18 | 28 |
|  | Final Total | 1,380 | 1,397 | 30,586 |
| Edges | Duration [min] | 207 | 140 | 410 |
|  | Local Thresholds | 9 | 0 | 1 |
|  | Selections | 145 | 66 | 96 |
|  | Additions | 152 | 63 | 557 |
|  | Final Total | 1,193 | 1,323 | 31,523 |

### Example anatomical statistics

To demonstrate its statistical power, the SLAVV software was used to automatically vectorize and manually classify a large volume of capillaries, venules, and arterioles. Figure 4A shows a 3D rendering of the vectorized vessels (color-coded by strand) beneath a maximum intensity projection of the superficial layers of the tiled (see Large volume tiling) Image 3. The vectorized vessels were idealized as collections of cylinders attached at the centers of their faces, and connected face to face (Figure 4B). Figure 4C shows lateral-area weighted histograms of vessel statistics (see Statistical analysis of vectors): depth, radius, and inclination (i.e. angle from *xy* plane). Figure 4D shows bulk statistics (length, area, volume, radius, and inclination) binned by depth (*z* coordinate). The total summary statistics (see Statistical analysis of vectors) and a binary mask (see Visualizations: Three-dimensional scalar fields) derived from this vector set serve as ground truth statistics and image segmentation for the automated SLAVV performance evaluation.

### Objective performance evaluation of fully-automated SLAVV

To remove the dependence on (subjective) manual annotation in determining ground truth images, two-photon images were simulated (see Simulating 2PM images) from an assumed ground truth anatomy with various levels of contrast and noise. To enable the tracking of bulk network statistic accuracy in addition to image segmentation accuracy, a ground truth vector set (with known network statistics) was used to generate the ground truth segmented image. Image segmentation performance was evaluated using ROC curves to show the voxel-by-voxel segmentation strength of the energy feature of vectors. Errors in bulk statistics were used as performance metrics because they additionally capture topographical and connectivity accuracy. Seven simulated images of varying quality were generated from the vector set from Image 3 (Table 1). The noise and contrast settings for the simulated images are shown in Figure 5A, and maximum intensity projections are shown in the top row of Figure 5B.

**Fig. 5.**
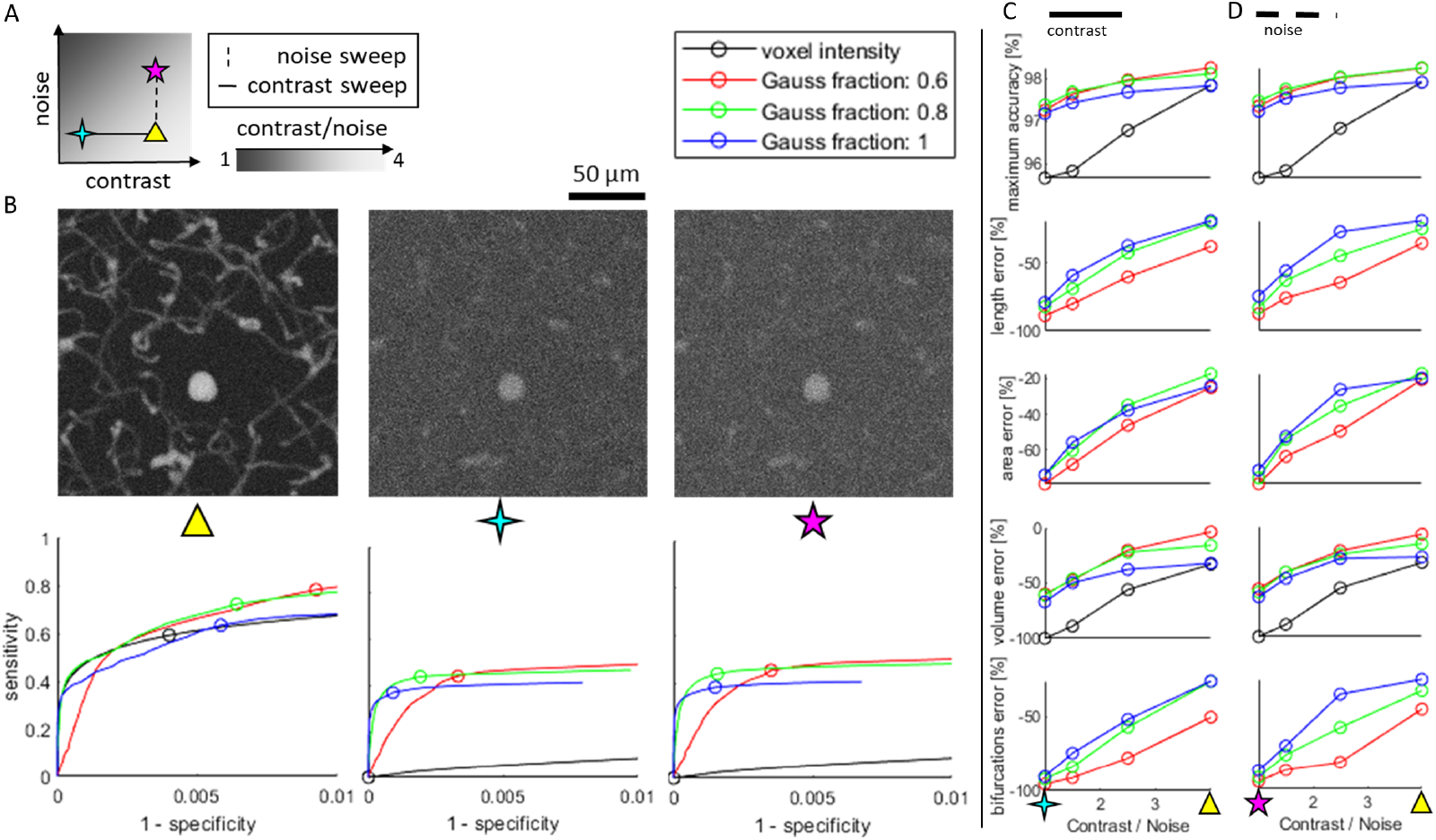
Objective performance of fully-automated SLAVV software. A. Simulated images of varying quality are generated from the vector set from Image 3 shown in Figure 4. Image quality is swept along contrast and noise axes, independently. Example maximum intensity projections are shown for three extremes of image quality (triangle: best quality, 4-point star: high noise, 5-point star: lowest contrast). The legend shows the labels for the four segmentation methods used in B-D. Images are vectorized using SLAVV with different amounts of Gaussian filter, *f_G_* (60, 80, or 100 % of matched filter length). B. Vasculature is segmented from three simulated images using four automated approaches: thresholding either voxel intensity or maximum energy feature on edge objects produced by three automated vectorizations. Voxel-by-voxel classification strengths of thresholded vectorized objects or voxel intensities are shown as ROC curves for three of the seven input images. Note that the ROC curves for the energy feature of vectors do not have support for every voxel, because not every voxel is contained in an extracted vector volume. Operating points with maximal classification accuracy are indicated by circles in the bottom row of B and plotted in the top row of C&D across all input images. C&D. Bulk network statistics (length, area, volume, and number of bifurcations) were extracted from vectors or binary images resulting from maximal accuracy operating points. Performance metrics were plotted against CNR (image quality) for a (C) contrast or (D) noise sweep. Thresholding vectorized objects to segment vasculature demonstrated a greater robustness to image quality than thresholding voxel intensities. Surface area, length, and number of bifurcations were not extracted from binary images, because these images were topologically very inaccurate.

The simulated images were vectorized to characterize the performance of the fully-automated (see Automated energy thresholding) SLAVV software. Three values (60, 80, and 100 %) were used for the *f_G_* processing parameter in the energy calculation step. Vertices were extracted for each energy image and classified using global thresholding with a dense sweep of thresholds. The curated vertex sets were rendered at the resolution of the input image and compared to the ground truth image voxel-by-voxel to calculate the confusion table enumerating the false positives *FP*, true positives *TP*, false negatives *FN*, and true negatives *TN*. The vertex set yielding the highest classification accuracy (*TP* + *TN*)*/*(*TP* + *TN* + *FP* + *FN*) was passed to the edge extraction step.

Edges were extracted for each vertex set and similarly classified using many global thresholds. Edge sets were rendered and voxel-by-voxel compared to the ground truth to find the sensitivity and specificity for each threshold. An ROC curve was made to observe the voxel-by-voxel classification strength of global thresholding on the edge objects (bottom row of Figure 5B). The edge set from each threshold sweep with maximal voxel-by-voxel classification accuracy was passed on to the network calculation step to compute statistics and compare to those of the ground truth.

The simulated image quality was benchmarked using a commonly-used, voxel-by-voxel, intensity thresholding classifier (Figure 5, black series) that has perfect accuracy in the limit of perfect image quality (CNR → ∞). To demonstrate the sensitivity of the intensity thresholding classifier to image quality, the volume was computed from the intensity thresholded image with the best voxel-by-voxel classification accuracy. For the higher quality images, the volume accuracy was comparable to that of the vectorized approaches, however, the topology was unconstrained, leading to salt and pepper segmentation errors. These topological errors made surface area calculation highly sensitive to noise for the intensity classifier approach (result not shown). For the lowest quality image, the intensity classifier yielded a maximal segmentation accuracy of around 95 %, corresponding to the operating point that assigns all voxels to background (0 % volume accuracy).

As a vessel segmentation tool on input images of poor quality (1 ≤ CNR ≤ 4), three fully-automated implementations of SLAVV outperformed the intensity classifier (Figure 5C and D. The peak, voxel-by-voxel classification accuracy of fully-automated SLAVV was above 97 % for all image qualities tested. However, the number of bifurcations detected was very sensitive to noise. A larger fraction of Gaussian in the filtering kernel (*f_G_ →*100 %) provided greater robustness to noise for bifurcation detection, but resulted in an underestimate of volume even for the higher quality input images.

## Discussion

### Advantages of the direct vectorization approach

The SLAVV method is advantageous because it is robust to input signal shape, quality, and resolution. Through efficient linear filtering, vessel centerlines and sizes are estimated along with an “energy” or goodness of fit metric. Vessel objects are extracted using simple algorithms to utilize the energy information along with size and topological (connectivity) constraints. Unlike image segmentation approaches, which classify voxels as true or false from grayscale images, the SLAVV approach directly extracts elementary vectors from the grayscale image. In doing so, it removes the needs for pre or post-image processing (interpolation, morphological filtering, skeletonization), which improves computational efficiency and removes sensitivities to image quality, resolution, and signal shape (spherical vs annular). Additionally, because the method is developed from first principles in signal processing, there is no machine learning required, and there is no need to train the software or create separate training and testing datasets. Although the SLAVV method does not segment the image before extracting vectors, it can nonetheless be used as a segmentation tool (by rendering the extracted vectors at the resolution of the original). Unlike voxel-by-voxel image segmentation methods, the output of SLAVV has guaranteed shape and connectivity constraints that are realistic for vascular networks.

### Performance of SLAVV on endothelial-labeled inputs

An endothelial labeled image, such as Image 1 is very difficult to segment using voxel-by-voxel classification, because the signal only outlines the object instead of filling its volume. The SLAVV approach is robust to this input signal shape, because it can be tuned to detect a combination of spherical and annular shapes. However, manual vector classification is less efficient on the endothelial-labeled Image 1 than the plasma-labeled Image 2 (Table 5), because the automated vector extraction is less robust to noise for the endothelial signal which is weaker and spans fewer voxels. Despite these obstacles, the SLAVV method enables the calculation of morphological statistics from endothelial contrast (Table 4) that would otherwise be very difficult. Additionally, because the initial filtering is linear, the superposition principle guarantees good performance on any combination of plasma- and endothelial- labeled inputs (result not shown).

### Anatomical statistic accuracy

The surface area density was observed to be uniform in depth (Figure 4C or D), consistent with an assumption of homogeneous chemical transport demand. Previous anatomical statistics describing cerebral microvasculature in mice have largely come from two-photon images of post-mortem brain tissue. In a comprehensive review by [10], the vascular volume density was reported to be between 0.5 and 1 % and the length density between 0.7 and 1.1 m/mm^3^. These results are comparable to the bulk statistics obtained in the cursory anatomical study shown in Figure 4 and Table 4. For example, the vectors extracted from Image 3 had a length density of 0.6 m/mm^3^. Interestingly, the corresponding volume density was 6 %. This larger volume estimate could be attributed to a higher vascular perfusion under anesthesia and in vivo compared to post-mortem imaging.

### Fully-automated SLAVV performance

The objective performance evaluation demonstrates a rigorous framework for testing the accuracy of vectorization techniques on a simulated image with known quality and ground truth. The ability of SLAVV to extract bulk network statistics such as surface area, length, and number of bifurcations is shown to be more robust to image quality than the voxel intensity classifier benchmark. The vectorizations utilized size, shape, and topographical information to achieve a greater robustness to image quality. Therefore, it is unsurprising that there was an image quality threshold below which the vector energy classifier outperformed the voxel intensity classifier.

### Future work: neurovascular plasticity

The vectorization software will be expanded to accept time-lapse images, automatically register vascular objects between imaging sessions, and manually curate any changes. Time-lapse images will be analyzed to estimate vascular plasticity statistics such as angiogenesis or angionecrosis. Time-resolved neurovascular statistics in murine cortex will be estimated with high accuracy and precision and in large volume. Good estimates of these statistics will inform fundamental neurovascular research. Measured statistics will be compared between treatment groups in a relevant medical experiment, for example to study the effect of a drug on capillary plasticity in adult murine cortex.

## Conclusion

We present the SLAVV method to vectorize microvascular networks directly from unprocessed, in vivo 2PM images of the mouse cortex. This vectorization method removes any preprocessing requirements (image segmentation, interpolation, denoising) by utilizing linear filtering and low-complexity vector extraction algorithms. Furthermore, there is no machine learning required, so the user does not need to generate separate training and testing datasets. The SLAVV method is shown to perform similarly on plasma- and endothelial-labeled images, enabling statistical calculations for the bulk network (length, area, volume, bifurcation frequency) and for individual capillaries, arterioles, or venules. Automated and manual processing costs are shown to scale proportionally with input image volume. Fully-automated SLAVV performance is proven robust to image quality compared to a common intensity-based thresholding approach. The SLAVV method is expected to enable longitudinal capillary tracking through accurate and efficient vectorization of time-lapse, in vivo images of mouse cerebral microvasculature.

## Acknowledgments

National Institutes of Health (NIH) (EB011556, NS082518, NS108484, T32EB007507)

